# Detecting cryptic ghost lineage introgression in four-taxon genomic datasets

**DOI:** 10.1101/2025.04.28.651118

**Authors:** Evan S Forsythe, Blaine S Pappa, Darren A Clavette, Devin Y Mendoza

**Affiliations:** Oregon State University-Cascades, Biology Program; Oregon State University, Department of Integrative Biology

## Abstract

**Premise:** Hybridization and introgression are pervasive evolutionary forces that have played fundamental roles in shaping the diversity of wild and domesticated plants. Four-taxon tests for introgression provide a reliable framework for detecting signatures of ancient introgression from genomic data, which have played an important role in revealing the reticulate nature of plant evolution; however, there is emerging evidence that a cryptic process known as ghost lineage introgression has the potential to dramatically skew interpretations of four-taxon introgression statistics, particularly our ability to determine the lineages involved in introgression. This ambiguity limits our ability to resolve the mechanisms and functional implications of introgression because it means we can determine neither the donor nor the recipient of introgressed alleles with confidence.

**Methods:** Here, we develop *ghostbuster*, a statistical test designed to detect ghost lineage introgression in genomic data based on patterns of sequence divergence. We employ coalescent simulations to test our method and ascertain the conditions under which it accurately identifies ingroup versus ghost lineage introgression. Finally, to demonstrate the utility of *ghostbuster* we apply it to a previously identified introgression event in the plant family Brassicaceae.

**Result:** Our simulations reveal that *ghostbuster* accurately distinguishes ghost lineage introgression from ingroup introgression across a wide range of introgression scenarios, with errors arising only when divergence events are closely spaced. Our analysis of empirical plant data reveals that the previously identified introgression likely constitutes ghost lineage introgression and, thus, was previously misinterpreted.

**Discussion:** Our analyses of simulated and empirical data demonstrate that *ghostbuster* will be a helpful tool in resolving reticulate evolution in plants and other taxa. We demonstrate the biological insights that *ghostbuster* provides by presenting an updated model of ghost lineage introgression in Brassicaceae, impacting our understanding of the molecular evolution of crop and model species in this important plant lineage. *Ghostbuster* code is freely available at: https://github.com/EvanForsythe/Ghost_introgression.

## Introduction

Hybridization is widespread across eukaryotes (Taylor and Larson 2019) and is especially prominent in plants (Mallet 2005; Yakimowski and Rieseberg 2014; Mallet et al. 2016). Hybridization, followed by backcrossing, leads to introgression of alleles between divergent species or populations, a phenomenon which has been well-documented in plants (Rieseberg and Soltis 1991; Rieseberg 2006; Suarez-Gonzalez et al. 2016; Forsythe et al. 2020a) and animals (Dasmahapatra et al. 2012), including humans (Green et al. 2010; Prüfer et al. 2014; Kuhlwilm et al. 2016). Advances in genome sequencing technology have led to the development of statistical approaches to detecting patterns of historical introgression in extant species (Green et al. 2010; Durand et al. 2011; Martin et al. 2015; Pease and Hahn 2015; Hahn and Hibbins 2019; Hibbins and Hahn 2021). Many of the most widely used statistics employ a four-taxon sampling approach to detect an abundance of derived alleles shared by non-sister species by detecting imbalanced site pattern or gene tree topology counts (Green et al. 2010; Durand et al. 2011; Martin et al. 2015; Martin and Jiggins 2017; Zheng and Janke 2018; Dagilis et al. 2021). Four-taxon statistics have been used to identify foundational cases of adaptive introgression (Dasmahapatra et al. 2012), extensive introgression impacting large portions of the genome (Fontaine et al. 2015; Forsythe et al. 2020a), and introgression events with profound significance to human health and medicine (Prüfer et al. 2014). Given the impact of introgression on the molecular evolution of important taxa, it is paramount to fully resolve a detailed understanding of these introgression events.

One of the most fundamental details regarding an introgression event is to resolve the lineages that exchanged alleles during historical gene flow (Hibbins and Hahn 2019, 2021; Forsythe et al. 2020b). Four-taxon tests are designed to reveal two alternative introgression events between two alternative pairs of non-sister taxa within the ingroup clade of the test species (Durand et al. 2011; Tricou et al. 2022b). These two alternative introgression scenarios are distinguished by the sign (positive versus negative) of the test statistic. Once the two species involved in an introgression event have been identified with a four-taxon test, auxiliary tests (Hibbins and Hahn 2019, 2021; Forsythe et al. 2020b), which make use of sequence divergence information, can be applied to resolve the directionality of introgression (i.e. the donor and recipient lineage). Furthermore, modified four-taxon tests have also been developed to identify the specific genomic regions (Martin et al. 2015; Martin and Jiggins 2017) that were introgressed between species, revealing the genomic and functional impacts of introgression.

Combining four-taxon site-pattern tests and divergence-based tests hold potential to resolve introgression events within sampled ingroup species; however, there is a growing body of evidence that suggests that interpretation of these tests is dramatically impacted by cryptic introgression stemming from unsampled or extinct lineages that diverged prior to the divergence of the sampled ingroup species (Hibbins and Hahn; Durand et al. 2011; Tricou et al. 2022a, 2022b; Tiley et al. 2023; Pang and Zhang 2024). Introgression from these so-called “ghost lineages” can skew allele patterns and gene tree topologies in a manner that exactly mimics the signature of ingroup introgression produced by four-taxon tests (Tricou et al. 2022b). This caveat is especially problematic because the implication of a misidentified ghost lineage introgression event means that neither the donor lineage nor the recipient lineage are correctly inferred (Tricou et al. 2022a; Pang and Zhang 2024). Considering that the vast majority of species that have existed on earth are either extinct or undescribed (Mora et al. 2011) and that fossil and archaic DNA evidence has demonstrated gene flow between now-extinct species (Slon et al. 2018), it is likely that at least some of the previously identified introgression events detected with four-taxon tests have been misinterpreted.

Given the potential for misinterpretation of introgression results, tools to help differentiate between ingroup and ghost lineage introgression are of value. Recent work thoroughly investigated the utility of existing heuristic methods (i.e. methods that use topology counts and genome-wide summary statistics) in accurately identifying ingroup versus ghost lineage introgression, revealing that heuristic methods perform poorly in their ability to correctly identifying ghost lineage introgression (Pang and Zhang 2024). In contrast, this study also demonstrated that a model-based approach, *Bayesian Phylogenetics and Phylogeography (BPP)* (Yang 2015; Flouri et al. 2020; Jiao et al. 2021) was able to accurately detect ghost lineage introgression. This study highlighted the utility of full likelihood demographic methods, which make use of both topology and sequence divergence information in sequence data (Hibbins and Hahn 2021), in accurately identifying modes of introgression. Based on these promising results, *BPP* is likely to become a valuable addition to introgression analysis workflows. However, the authors also note that such methods are computationally expensive compared to analogous heuristic-based methods, presenting a challenge for analyzing large genomic datasets (Pang and Zhang 2024). In addition to computational cost, model-based methods may be prone to error associated with inaccurate parameter estimation and model overfitting (White et al. 2016). Given these limitations, a heuristic method that explicitly tests for ingroup versus ghost lineage introgression would be a valuable addition to analytical workflows.

Here, we present *ghostbuster*, a novel statistical test to distinguish ingroup from ghost lineage introgression in four-taxon genomic data. *Ghostbuster* provides an efficient heuristic approach to using sequence divergence summary statistics to test introgression hypotheses. We define the expected divergence profiles under ingroup versus ghost lineage introgression and perform multispecies coalescent simulations to test the performance and accuracy of *ghostbuster* under different introgression scenarios. Finally, we apply *ghostbuster* to a previously described introgression event in the plant family Brassicaceae and present a revised model for introgression in the group.

## Methods

### Implementation of the ghostbuster test

*Ghostbuster* is designed to analyze the same data used for four-taxon introgression tests (Green et al. 2010; Durand et al. 2011) and is intended for use only after a significant signature of introgression has already been detected. *Ghostbuster* requires fasta input files with DNA sequences for single-copy genes/loci/windows spanning the genome. Each file must contain a sequence from at least four species (P1, P2, P3, and an outgroup) (Fig. 1) used in upstream four-taxon tests. Sequences from extra species are pruned from files prior to multisequence alignment.

**Fig. 1:**
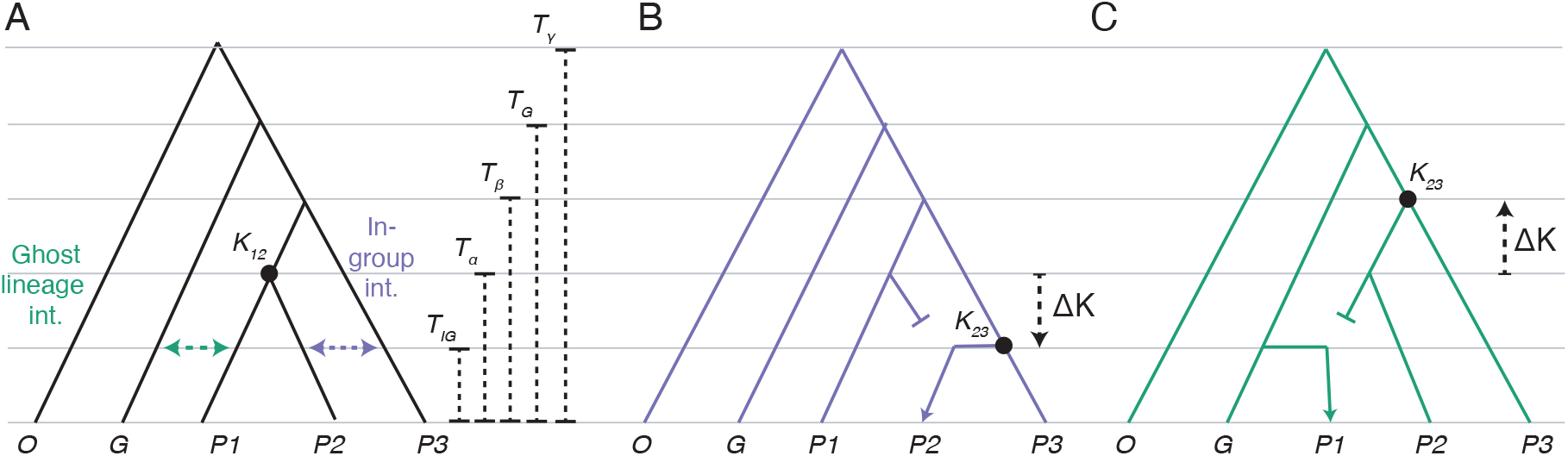
Expected sequence divergence under ingroup versus ghost lineage introgression models. (**A**) Species tree (black branches) with possible introgression events (green and purple arrows) and tree depth times for speciation and introgression events (dotted lines). The K_12_ node indicates the node of interest on gene trees that exhibit the species topology [(P1,P2),P3)]. (**B-C**) Gene tree resulting from ingroup (**B**) and ghost lineage (**C**) introgression. The K_23_ node indicates the node of interest on gene trees that exhibit the introgression topology [(P2,P3),P1)], displayed in both introgression scenarios. ΔK symbols and arrows indicate the expected difference between K_12_ and K_23_ when comparing introgressed genes to non-introgressed genes.

For each input file, the following steps are performed: multiple sequence alignment with *MAFFT* (Katoh and Standley 2013), maximum likelihood inference with *IQ-TREE* (Nguyen et al. 2015), which employs *ModelFinder* (Kalyaanamoorthy et al. 2017), and finally, topology and branch length analyses using *biopython* (Chapman and Chang 2000) tools (Fig. 2). Each tree is rooted by the user-specified outgroup and branch lengths (measured in substitution per site) are calculated from each gene tree after topology is assessed. A divergence value for each node is obtained by summing the branch lengths of the two external branches subtending the node.

**Fig. 2:**
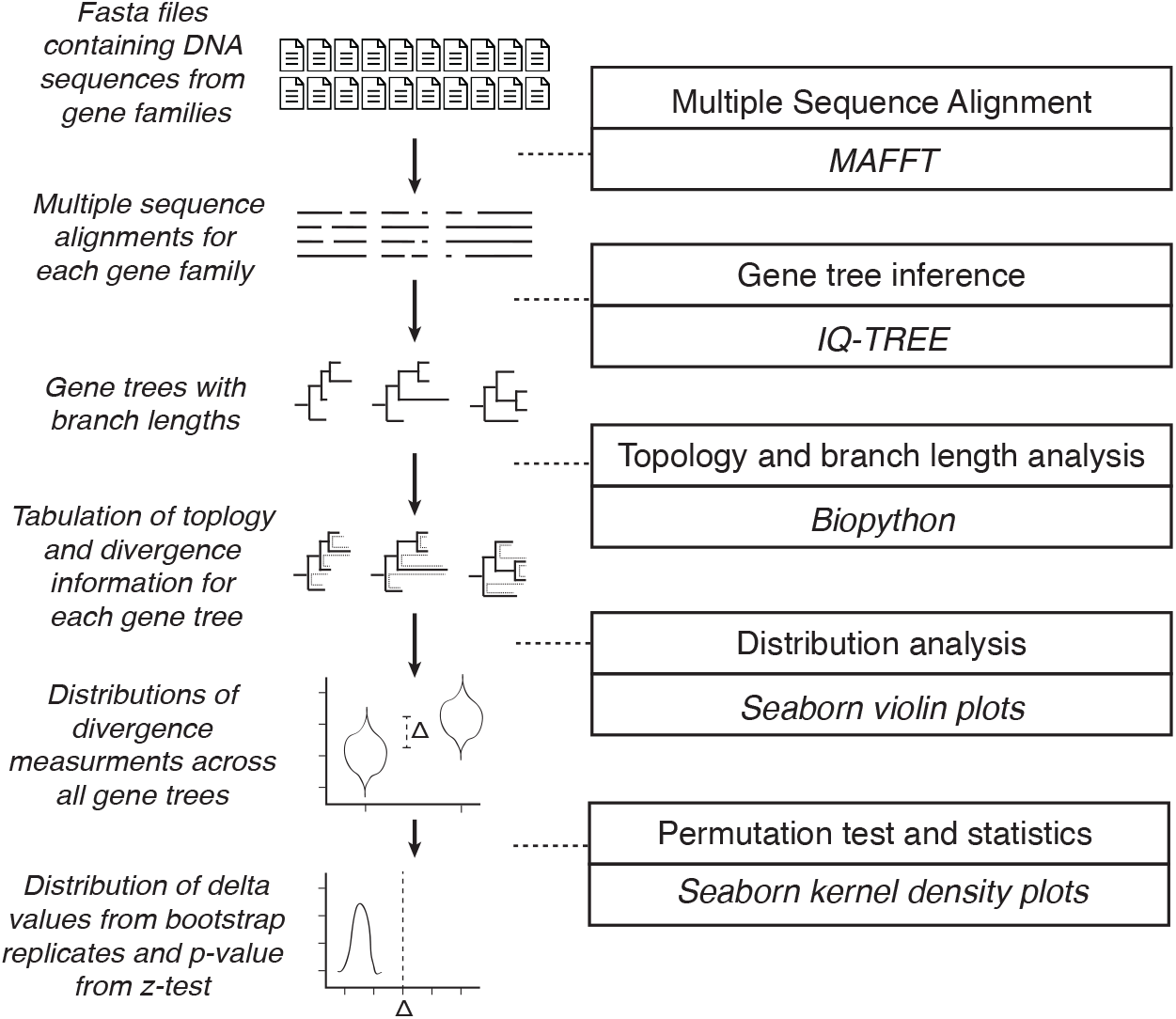
Analytical workflow of the ghostbuster test. Major steps of the *ghostbuster* test, implemented as a python tool available at https://github.com/EvanForsythe/Ghost_introgression.

*Ghostbuster* assumes that the user-specified species tree topology is (((P1,P2)P3)O), and calculates node depth K_12_ from gene trees exhibiting this topology. Gene trees with the topology (((P2,P3)P1)O) are used to calculate K_23_. Gene trees with the topology (((P1,P3)P2)O) are presumed to originate from incomplete lineage sorting (see Discussion) and are not used by *ghostbuster*. ΔK is calculated by subtracting the average from all K_23_ values from the average from all K_12_ values (see Eq.1). To assess the significance of the ΔK, *ghostbuster* performs a bootstrap replication permutation test with 100 replicates. For each bootstrap replicate, gene trees are randomly sampled (with replacement) and K_12_, K_23_ are estimated for the sample to produce a resampling distribution of ΔK. Similar to the approach used by (Forsythe et al. 2020b), a two-sided z-test is performed to assess whether the distribution significantly differs from zero.

### DNA sequence simulations of ingroup and ghost lineage introgression scenarios

To test *ghostbuster* performance, we developed simulations using *tskit* (https://tskit.dev/) and *msprime* (Baumdicker et al. 2022) multispecies coalescence simulation software packages. We implemented simulations using the script, *Data_simulations*.*py*, available with *ghostbusters* (https://github.com/EvanForsythe/Ghost_introgression). Speciation events were simulated via the *add_mass_migration()* command with the proportion argument set to 1.0. In a reverse-time (coalescent) perspective, these events are akin to two populations merging, however, when considered from a forward-time direction, they are akin to a speciation event. Introgression events were simulated with the *add_mass_migration()* command with the proportion argument set to 0.2, simulating (in forward-time) an introgression event in which 20% of alleles are introgressed. Introgression was simulated as either ‘ghost’ or ‘true’ by setting the “destination” (i.e. the source in forward-time) of migration. In all cases, we simulate the presence of a ghost lineage, but we never include the resulting sequence from this Ghost taxon in downstream analyses, thus effectively mimicking an ‘unknown’ ghost lineage.

Our default time points (in coalescent units) were set to the following: T_IG_=40,000; T_α_=80,000; T_β_ = 120,000; T_G_=160,000; T_γ_=200,000 (Fig 1A). For parameter scan analyses, we multiplied all time point (Fig. 4A) or the T_IG_ time point (Fig. 4B) by a scaling factor (SF).

**Fig. 3:**
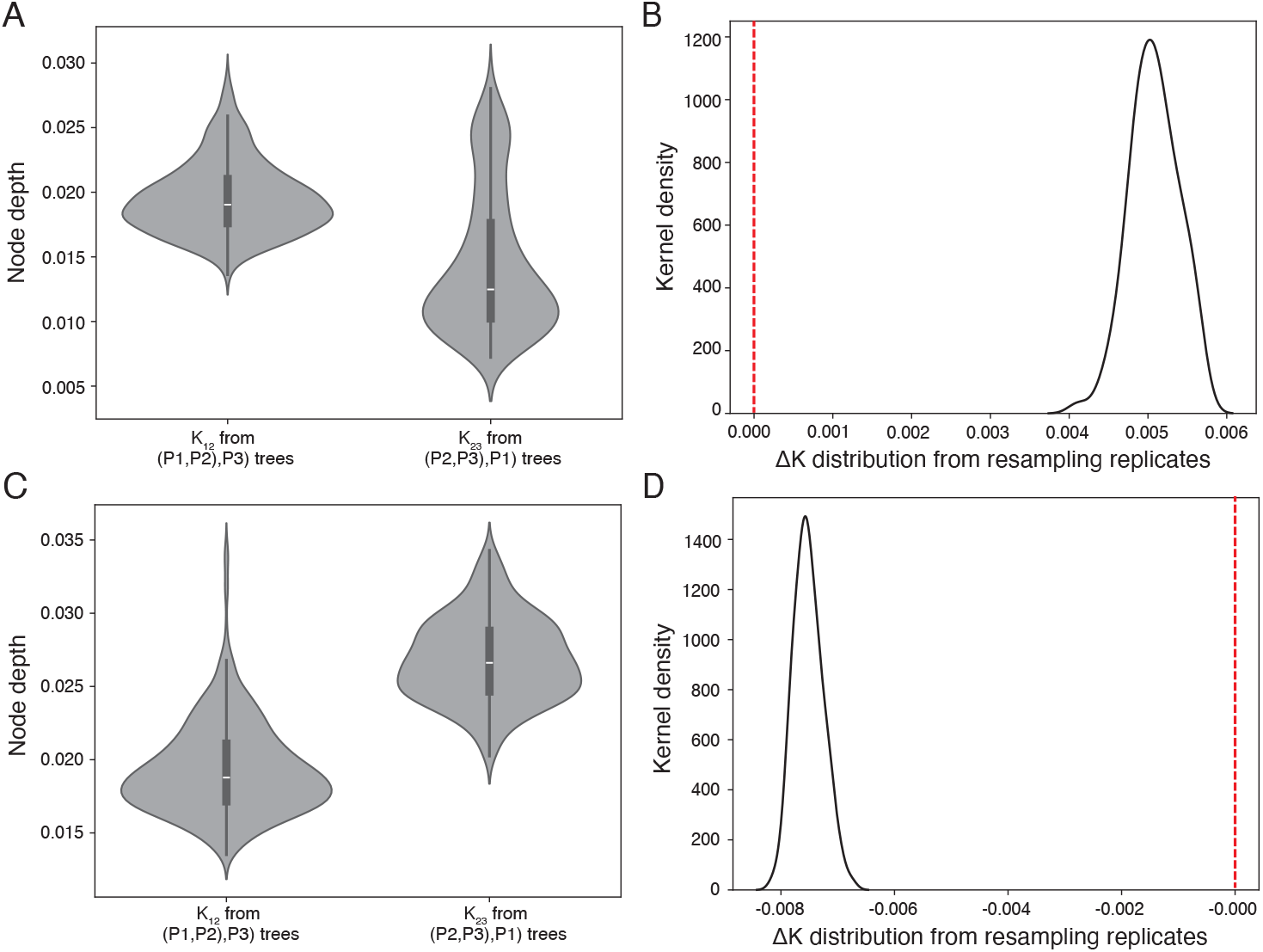
Applying the *ghostbuster* test to simulated data. *Ghostbuster* output on simulated “ingroup” introgression dataset in which introgression was simulated from P3 to P2 (**A-B**) or simulated “ghost lineage” introgression, in which introgression was simulated from a ghost lineage into P1 (**C-D**). (**A and C**) The distribution of node depths obtained from gene trees with the speciation topology or the introgression topology. (**B and D**) shows ΔK distribution from 100 bootstrap resampling replicates. Both distributions significantly (p<<0.01) differ from zero, indicating significant signatures of ingroup (**B**) and ghost lineage (**D**) introgression.

**Fig. 4:**
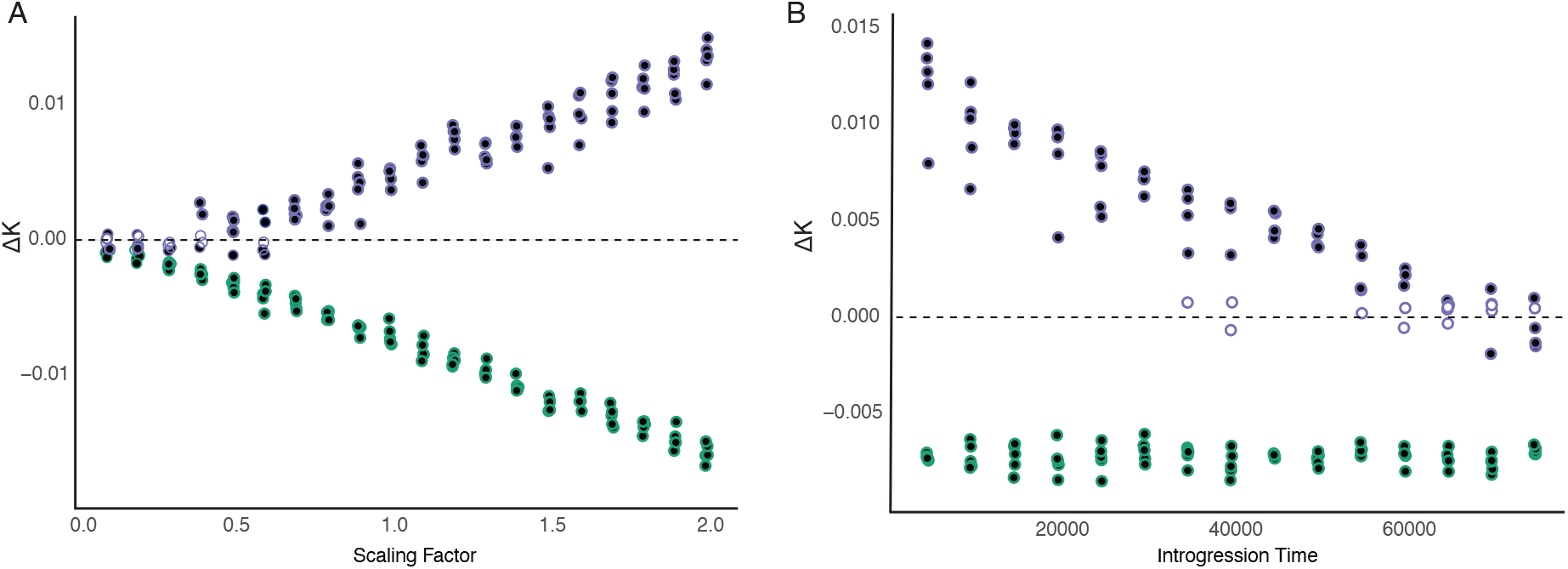
Testing the robustness of the *ghostbuster* method under simulated introgression scenarios. (**A-B**) ΔK measurements from *ghostbuster* runs on datasets with simulated ingroup (purple points) and ghost lineage (green points) introgression. Positive and negative ΔK measurements from the *ghostbuster* test indicate ingroup and ghost lineage introgression, respectively. (**A**) Parameter scan of varying scaling factor values used to scale the simulated timing of all divergence and reticulation events proportionally. Larger numbers equate to larger tree heights on simulated trees. (**B**) Parameter scan of introgression scenarios in which introgression time is changed while all other time points are held constant. Larger values indicate introgression (T_IG_) occurring at deeper time points approaching the timing of the speciation event (T_α_). Filled points indicate significant (z-test; *p*<0.05) *ghostbuster* test result. Open points indicate that ΔK distributions did not significantly differ from zero.

Additional parameters used during sequence simulations were: sequence length=10,000,000 bp; mutation rate=0.0000001 (mutations per bp per generation), recombination rate=0.000000001 (recombination events per bp per generation); and Ne=10,000. These parameters were held constant for the simulations in this study.

The output of the above simulations is a ‘full chromosome’ alignment comprising numerous introgressed and non-introgressed haplotype blocks. From this full chromosome alignment, we achieved single-gene fasta files for input into *ghostbuster* by splitting the full chromosome alignment into 1,000 non-overlapping windows, which were used to infer “gene trees” in downstream *ghostbuster* analyses.

### Testing for ghost lineage introgression in empirical plant genomic data

To test *ghostbuster* on an empirical dataset, we used genomic data from a previous study that identified introgression between model species in the Brassicaceae family (Forsythe et al. 2020a). We obtained coding sequence (CDS) alignments for single-copy genes used in the prior study. We applied the *ghostbuster* test to a representative species from each clade using the species tree topology shown in Figure 5. Analyses were performed on a *Linux* server equipped with dual AMD EPYC 7713 processors using four parallel threads. Analyses were repeated on a Macintosh laptop, yielding similar runtimes and results.

**Fig. 5:**
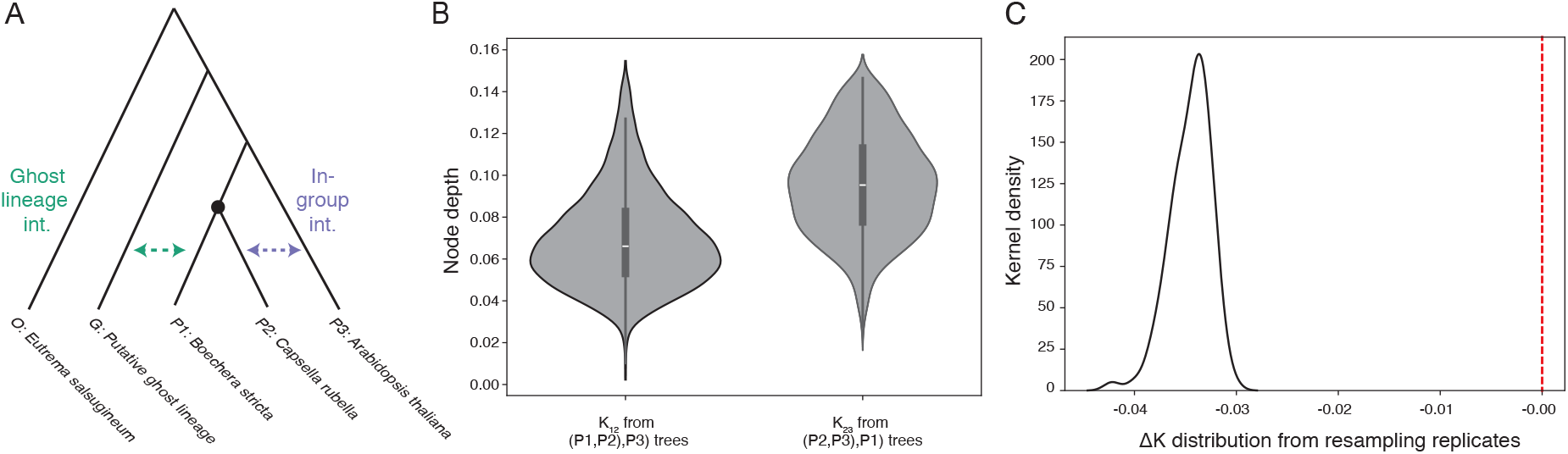
Testing for ghost lineage introgression in a previously identified introgression event in Brassicaceae. (**A**) Species relationships among test species as indicated by the major topology from all gene trees. Green and purple arrows indicate alternative introgression scenarios that would conceivably result in the previously observed (P2,P3),P1) topology gene trees. (**B**) The distribution of node depths obtained from gene trees with the speciation topology (left) or the introgression topology (right). (**C**) ΔK distribution from 100 bootstrap resampling replicates.

## Results

### Expected sequence divergence under ingroup and ghost lineage introgression

Widely used four-taxon tests for introgression rely on the topology of gene trees or corresponding site-patterns to indicate the presence of introgression. However, ingroup and ghost lineage introgression can lead to identical topologies and site patterns (Tricou et al. 2022a, 2022b), meaning these pieces of evidence cannot differentiate between alternative introgression scenarios. We reasoned that the divergence profiles of introgressed and non-introgressed loci could help resolve the species involved in introgression events. To outline our expectations for introgressed gene trees, we defined time points, corresponding to introgression events (T_IG_) and speciation events (T_α_, T_β_, T_G_, and T_γ_) in a four-taxon (not including a ghost lineage) introgression scenario (Fig. 1A). Phylogenetic estimates of the relative timing of divergence events are obtainable by calculating sequence divergence (i.e. node depth) between pairs of taxa on gene trees (Fig 1). Gene trees that were not introgressed are expected to display the ‘species topology’ and will inherently contain a node representing the divergence of P1 and P2; the node depth of this node (K_12_) should always correspond to T_α_ (Fig. 1A). On the other hand, gene trees that underwent introgression will display the ‘introgressed topology’ and, as such, will inherently contain a node representing the divergence of P2 and P3 with node depth K_23_ (Fig. 1B-C).

Importantly, unlike K_12_, K_23_ will differ depending on whether introgression is ingroup (K_23_ = T_IG_) (Fig. 1B) versus ghost lineage (K_23_ = T_β_) introgression (Fig. 1C).

To detect sequence divergence profile of introgression events, we propose a simple heuristic, the ΔK statistic:

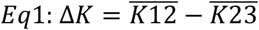

Under ingroup introgression, we expect ΔK to be positive, such that:

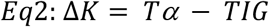

Under ghost lineage introgression, we expect ΔK to be negative, such that:

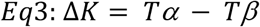

Our ΔK statistic provides a measurable readout of the divergence profile of introgression events that are detected with four-taxon statistics.

### Ghostbuster: a divergence-based four-taxon test for ghost lineage introgression

We created a statistical framework and analytical pipeline for asking whether divergence profiles are most consistent with ingroup versus ghost lineage introgression (Fig. 2). Our workflow takes multiple DNA sequence files as input, each of which represents a gene, locus, or genomic window (hereafter, “gene”) from the four test species suspected to have undergone introgression. For each gene, we perform multiple sequence alignment, gene tree inference, topology and branch length analysis to extract the node depth estimates described above and use genome-wide averages to estimate ΔK for the dataset. To assess whether our ΔK point estimate is significantly positive or negative, we employ a bootstrap resampling approach (Forsythe et al. 2020b) and use the resulting distribution of resampled ΔK estimates to perform a z-test. We implemented *ghostbuster* as a python tool that requires a minimal set of dependencies and computational power. Our approach provides a scalable framework for testing the divergence profile of introgression events.

### Testing ghostbuster performance on simulated genomes

To test our *ghostbuster* approach, we simulated DNA sequence evolution for populations described in Fig 1A. Ingroup introgression was achieved by simulated gene flow from P3 to P2. Ghost lineage introgression was achieved by simulating gene flow from the ghost lineage (G) into P1. This approach mimics ghost lineage introgression because all downstream analyses omit the ghost population, only making use of O, P1, P2, P3 in divergence analyses (see Methods).

To demonstrate the output of ghostbuster, we applied the test to genomes simulated to have undergone ingroup introgression (Fig. 3A-B) and ghost lineage introgression (Fig. 3C-D) at our default timepoints (see Methods). Our results show that simulated ingroup introgression leads to a higher K_12_ distribution compared to the K_23_ distribution (Fig. 3A). This divergence profile resulted in a ΔK distribution significantly (p<0.001) greater than zero (Fig. 3B), consistent with our theoretical expectations for ingroup introgression. When we simulated ghost lineage introgression, we observed the opposite pattern for K_12_ and K_23_ distributions (Fig. 3C), resulting in a significantly negative (p<0.001) ΔK distribution (Fig. 3D), again consistent with our expectations. These results provide a proof-of-principle for the effectiveness of our *ghostbuster* test.

To evaluate robustness of the *ghostbuster* test, we performed parameter scans, in which we altered the timing of simulated speciation and introgression events. First, to explore the effect of overall tree height on ghostbuster results, we employed a ‘scaling factor’ (SF) and multiplied all time points by this number to achieve simulated genomes with overall larger (scaling factor >1) or smaller (scaling factor <1) levels of divergence (Fig 4A). We found that increased tree height provided increased resolution for the *ghostbuster* test. The effect-size of ΔK grew increasingly positive (ingroup introgression) or negative (ghost lineage introgression) as SF increased. We simulated five replicate genomes for each SF value. Ghostbuster significantly inferred the correct mode of introgression with 100% accuracy for all SF ≥ 0.7 runs. When applied to simulated ghost lineage introgression, *ghostbuster* inferred the correct mode for all replicates and scaling factor values, with the exception of a single insignificant result for one replicate at SF=0.2. In contrast, *ghostbuster* accuracy suffered when applied to simulated ingroup introgression on SF<0.7 runs, as evidenced by many cases of erroneous ΔK estimates below zero at low SF values. In some cases, these erroneous results were statistically significant, indicating that ghostbuster is prone to error under ‘rapid speciation’ scenarios, when divergence events occur in close succession.

To test the effect of the relative timing of speciation and introgression events, we also performed a parameter scan in which we varied only the timing of introgression (T_IG_), while leaving all other time points constant. For these simulations, increasing T_IG_ brings it closer to T_α_ (Fig. 4B). For ghost lineage introgression simulations, we found that ΔK values are unaffected by T_IG_. This is consistent with our theoretical expectations because ΔK is affected only by T_α_ and T_β_ in ghost lineage introgression scenarios (Eq. 3), which are unchanged in this analysis. In contrast, ingroup ΔK are heavily impacted by T_IG_, with increasing rate of insignificant and significantly erroneous results at high T_IG_ values. This result again highlights that *ghostbuster* accuracy suffers when divergence and introgression occur in short order. Taken together, our results show that *ghostbuster* is a reliable test under most introgression scenarios, but lacks resolution under rapid speciation (see Discussion).

### Applying ghostbuster to resolve introgression in Brassicaceae

With an understanding of *ghostbuster* performance on simulated data, we applied the *ghostbuster* test to a previously described introgression event in the plant family Brassicaceae (Forsythe et al. 2020a). We used the most frequent gene tree topology as our species topology (Fig. 5A). In the previous study, significant four-taxon test results were interpreted as evidence of ingroup introgression (Fig. 5A; purple arrow); however, the authors also discussed the possibility of a cryptic ghost lineage introgression event (Fig. 5A; green arrow) but lacked a statistical framework for testing these alternative hypotheses. To resolve this ambiguity, we applied *ghostbuster* to the same data. We observed a significantly negative (p<0.001) ΔK distribution, indicating that this previously described introgression event is likely a case of ghost lineage introgression (see Discussion). These results highlight the utility of *ghostbuster* in resolving introgression results, providing a new basis for interpreting four-taxon tests for introgression.

## Discussion

### A fast heuristic that explicitly tests for ghost lineage introgression

Our *ghostbuster* analysis of Brassicaceae alignments took 22 minutes to complete using four parallel threads for the phylogenetic inference steps. This runtime using only modest computational resources means that *ghostbuster* can accomplish real-life analyses using computing resources present in most modern base-model laptops. This efficiency is derived from the heuristic nature of comparing average node depths rather than traversing joint distributions of gene tree topologies and coalescent times. The most computationally intensive step of the *ghostbuster* workflow is the maximum likelihood gene tree inference step, which we achieve with *iqtree*. As part of gene tree and branch length inference, we employ *iqtree*’s “-m TEST” options to identify the best model of evolution for phylogenetic inference. Given model-based phylogentic inference step, it could be argued that *ghostbuster* is not strictly a heuristic method. However, the overall efficiency for the test and the incorporation of summary statistics make *ghostbuster* best aligned with site-pattern and gene tree topology-based methods that have been characterized as heuristic-based methods (Pang and Zhang 2024). In any case, *ghostbuster* shows promise as a fast and accurate test that can be routinely applied to the same four-taxon datasets that are widely used to indicate the presence of an introgression event.

### Reduced phylogenetic resolution during rapid speciation

Our simulation analyses sought to explore coalescence time parameters to understand the limits of *ghostbuster* performance (Fig. 4). These parameter scans revealed the *ghostbuster* test does not perform well under scenarios in which only a small amount of time separates the speciation event (T_α_) from the introgression event (T_IG_). Lack of phylogenetic resolution in these scenarios is not entirely surprising, given that the process of incomplete lineage sorting (ILS) is known to introduce alternative gene tree topologies with skewed coalescence times (Hibbins and Hahn 2021). Multispecies coalescent simulations inherently model the process of incomplete lineage sorting (i.e. deep coalescence), the extent of which is directly related to time between events (Hudson 2002). Consistent with the idea that ILS introduces noise to *ghostbuster*, the simulations that produced erroneous or insignificant *ghostbuster* results were also the simulations that produced the most gene trees with a topology in which P1 and P3 are sister (data not shown), which likely result from ILS.

Moreover, in addition to introducing a third gene tree topology on four-taxon trees, ILS is also expected to shift coalescent times, resulting in gene trees with deeper node depths. This impact affects gene trees displaying all three possible topologies, meaning that the genome-wide distribution of node depths across gene trees will be a mixed distribution of ILS trees and non-ILS trees. Consistent with this expectation, we observe a bimodal K_23_ distribution in our simulated ingroup introgression example (Fig. 3A). Despite ILS adding noise, the introgression signal (lower peak) vastly outweighed the ILS signal (higher peak), leading to a significant and accurate *ghostbuster* result. While cases in which ILS is more prominent should be interpreted with caution, we do not anticipate it posing a major barrier in the practical application of *ghostbuster*. This is because the conditions that would render *ghostbuster* ineffective are also likely to undermine upstream four-taxon introgression tests, making them less likely to produce a significant signature of introgression in the first place. As a result, a *ghostbuster* analysis would not be warranted under those conditions.

### An updated model of introgression in Brassicaceae

We previously interpreted four-taxon results in Brassicaceae as an indication of ingroup introgression (Forsythe et al. 2020a). However, our *ghostbuster* analysis indicates that this introgression event is likely introgression from an unknown ghost lineage (Fig. 5), indicating a novel alternative model for introgression in this important group of model and crop plants. We initially hypothesized the presence of this introgression event based on evidence of phylogenetic incongruence between nuclear and chloroplast gene trees (Beilstein et al. 2006, 2008) and subsequent genome-wide phylogenomic analysis revealed incongruence among nuclear genes, with ~88% of nuclear genes displaying a major topology and ~8% of nuclear genes displaying a minor topology (Forsythe et al. 2020a). The most parsimonious interpretation of these highly unbalanced ratios is that the major topology represents the ‘true species branching order’ and the minor topology represents introgressed genes. However, based on conflicting divergence values in our prior study, we explored an alternative hypothesis in which the minor topology represents the species branching order, while the major topology is the result of extensive introgression, affecting ~88% of nuclear genes. This interpretation was based on a similar pattern and interpretation of introgression in mosquitos (Fontaine et al. 2015). However, a later study reinterpreted the mosquito introgression data considering ghost lineage introgression and found that underlying assumptions about branch lengths do not hold true (Tricou et al. 2022a).

Consistent with this, our updated model suggests that the minor topology is indeed the result of ghost lineage introgression. However, it should be noted that the *ghostbuster* framework developed here does not explicitly test between different species trees and, therefore, cannot be used to rule-out the possibility of extensive introgression impacting most of the genome. Despite this, the growing understanding of ghost lineage introgression and its impacts on branch lengths favors our updated model as the most plausible scenario.

Our previous study of introgression in Brassicaceae found evidence that cytonuclear selection acting during introgression (Forsythe et al. 2020a). Based on previous interpretations (under the assumption of ingroup introgression), the recipient of this introgression was presumed to be ‘Clade C’, which includes the emerging biofuel crop, *Camelina sativa*. Our updated model of ghost lineage introgression suggests that, instead, the recipient of introgression is Clade B, represented by *Boechera stricta*, a genetic model used to study the genetics of asexual reproduction in plants (Schranz et al. 2006). It should be noted that our new model does not invalidate the prior findings related to cytonuclear selection, but it does change our understanding of the lineage in which this selection acted, which is an important component of resolving an introgression event. This updated interpretation of introgression highlights the biological insights gained from testing for ghost lineage introgression. Moreover, this new model, combined with the relative abundance of genomic resources available in the Brassicaceae family, creates an exciting opportunity to search for close relatives/descendants of the putative donor ghost lineage among extant Brassicaceae species.

Finally, given that we observed inconclusive and erroneous results in some scenarios in our simulated data (Fig. 4), it is important to ask whether Brassicaceae introgression falls in the range of divergence times for which *ghostbuster* performance is robust. Our best measure of this comes from comparing the node depth values in our simulated (Fig. 3) versus empirical (Fig. 5) data. This comparison shows that the node depths in the Brassicaceae analysis (Fig. 5B and C) fall on the high end of the spectrum of simulation scenarios we tested. Moreover, the shapes of the node depth distributions do not appear to be bimodal. Both observations suggest that our empirical analysis was not meaningfully impacted by ILS, which is an encouraging sign that our *ghostbuster* results for the Brassicaceae data are reliable. Taken together, our work adds important theoretical and methodological advancement to a growing body of work that seeks to resolve the fine details of introgression events identified via tractable four-taxon tests.

## Author Contributions

ESF conceived the project. ESF, DAC, and DYM wrote computer code for the analyses. BSP tested the code and ran analyses. ESF and BSP drafted and edited the manuscript.

## Acknowledgments

Thank you to Daniel Sloan and members of the Sloan laboratory for helpful discussion. This work was supported by funding from the National Science Foundation (IOS-2114641) awarded to ESF.

## Data Availability Statement

*Ghostbuster* code is freely available at: https://github.com/EvanForsythe/Ghost_introgression.

Python scripts used to generate all simulated data are also available with the *ghostbuster* distribution. Data used for empirical analyses in this work are available with the original publication.

## References

1. Baumdicker F, Bisschop G, Goldstein D, Gower G, Ragsdale AP, Tsambos G, Zhu S, Eldon B, Ellerman EC, Galloway JG, et al. Efficient ancestry and mutation simulation with msprime 1.0. Genetics. 2022:220(3). 10.1093/genetics/iyab229

2. Beilstein MA, Al-Shehbaz IA, and Kellogg EA. Brassicaceae phylogeny and trichome evolution. Am J Bot. 2006:93(4):607–619. 10.3732/ajb.93.4.607

3. Beilstein MA, Al-Shehbaz IA, Mathews S, and Kellogg EA. Brassicaceae phylogeny inferred from phytochrome A and ndhF sequence data: tribes and trichomes revisited. Am J Bot. 2008:95(10):1307–27. 10.3732/ajb.0800065

4. Chapman BA and Chang JT. Biopython: Python tools for computational biology. ACM SIGBIO Newsletter. 2000:20(August):15–19.

5. Dagilis AJ, Peede D, Coughlan JM, Jofre GI, D’Agostino ERR, Mavengere H, Tate AD, and Matute DR. 15 years of introgression studies: quantifying gene flow across Eukaryotes. 2021. 10.1101/2021.06.15.448399

6. Dasmahapatra KK, Walters JR, Briscoe AD, Davey JW, Whibley A, Nadeau NJ, Zimin A V., Hughes DST, Ferguson LC, Martin SH, et al. Butterfly genome reveals promiscuous exchange of mimicry adaptations among species. Nature. 2012:487(7405):94–98. 10.1038/nature11041

7. Durand EY, Patterson N, Reich D, and Slatkin M. Testing for Ancient Admixture between Closely Related Populations. Mol Biol Evol. 2011:28(8):2239–2252. 10.1093/molbev/msr048

8. Flouri T, Jiao X, Rannala B, and Yang Z. A bayesian implementation of the multispecies coalescent model with introgression for phylogenomic analysis. Mol Biol Evol. 2020:37(4):1211–1223. 10.1093/molbev/msz296

9. Fontaine MC, Pease JB, Steele A, Waterhouse RM, Neafsey DE, Sharakhov I V., Jiang X, Hall AB, Catteruccia F, Kakani E, et al. Extensive introgression in a malaria vector species complex revealed by phylogenomics. Science (1979). 2015:347(6217):1–6. 10.1126/science.1258522

10. Forsythe ES, Nelson ADL, and Beilstein MA. Biased gene retention in the face of introgression obscures species relationships. Genome Biol Evol. 2020a:12(9):1646–1663. 10.1093/GBE/EVAA149

11. Forsythe ES, Sloan DB, and Beilstein MA. Divergence-based introgression polarization. Genome Biol Evol. 2020b:12(4):463–478. 10.1093/gbe/evaa053

12. Green RE, Krause J, Briggs AW, Maricic T, Stenzel U, Kircher M, Patterson N, Li H, Zhai W, Fritz MH-Y, et al. A Draft Sequence of the Neandertal Genome. Science (1979). 2010:328(5979):710–722. 10.1126/science.1188021

13. Hahn MW and Hibbins MS. A Three-Sample Test for Introgression. Mol Biol Evol. 2019:36(12):2878–2882. 10.1093/molbev/msz178

14. Hibbins MS and Hahn MW. The timing and direction of introgression under the multispecies network coalescent. Genetics. 2019:211(March):1059–1073.

15. Hibbins MS and Hahn MW. Phylogenomic approaches to detecting and characterizing introgression. Genetics. 2021. 10.1093/genetics/iyab173

16. Hibbins MS and Hahn MW. Phylogenomic approaches to detecting and characterizing introgression. 1–42.

17. Hudson RR. Generating samples under a Wright-Fisher neutral model of genetic variation.

18. Jiao X, Flouri T, and Yang Z. Multispecies coalescent and its applications to infer species phylogenies and cross-species gene flow. Natl Sci Rev. 2021:8(12). 10.1093/nsr/nwab127

19. Kalyaanamoorthy S, Minh BQ, Wong TKF, Von Haeseler A, and Jermiin LS. ModelFinder: Fast model selection for accurate phylogenetic estimates. Nat Methods. 2017:14(6):587– 589. 10.1038/nmeth.4285

20. Katoh K and Standley DM. MAFFT multiple sequence alignment software version 7: Improvements in performance and usability. Mol Biol Evol. 2013:30(4):772–780. 10.1093/molbev/mst010

21. Kuhlwilm M, Gronau I, Hubisz MJ, De Filippo C, Prado-Martinez J, Kircher M, Fu Q, Burbano HA, Lalueza-Fox C, De La Rasilla M, et al. Ancient gene flow from early modern humans into Eastern Neanderthals. Nature. 2016:530(7591):429–433. 10.1038/nature16544

22. Mallet J. Hybridization as an invasion of the genome. Trends Ecol Evol. 2005:20(5):229–237. 10.1016/j.tree.2005.02.010

23. Mallet J, Besansky N, and Hahn MW. How reticulated are species? BioEssays. 2016:38(2):140–149. 10.1002/bies.201500149

24. Martin SH, Davey JW, and Jiggins CD. Evaluating the use of ABBA-BABA statistics to locate introgressed loci. Mol Biol Evol. 2015:32(1):244–257. 10.1093/molbev/msu269

25. Martin SH and Jiggins CD. Interpreting the genomic landscape of introgression. Curr Opin Genet Dev. 2017:47:69–74. 10.1016/j.gde.2017.08.007

26. Mora C, Tittensor DP, Adl S, Simpson AGB, and Worm B. How many species are there on earth and in the ocean? PLoS Biol. 2011:9(8). 10.1371/journal.pbio.1001127

27. Nguyen LT, Schmidt HA, Von Haeseler A, and Minh BQ. IQ-TREE: A fast and effective stochastic algorithm for estimating maximum-likelihood phylogenies. Mol Biol Evol. 2015:32(1):268–274. 10.1093/molbev/msu300

28. Pang XX and Zhang DY. Detection of Ghost Introgression Requires Exploiting Topological and Branch Length Information. Syst Biol. 2024:73(1):207–222. 10.1093/sysbio/syad077

29. Pease JB and Hahn MW. Detection and Polarization of Introgression in a Five-Taxon Phylogeny. Syst Biol. 2015:64(4):651–662. 10.1093/sysbio/syv023

30. Prüfer K, Racimo F, Patterson N, Jay F, Sankararaman S, Sawyer S, Heinze A, Renaud G, Sudmant PH, De Filippo C, et al. The complete genome sequence of a Neanderthal from the Altai Mountains. Nature. 2014:505(7481):43–49. 10.1038/nature12886

31. Rieseberg LH. Hybrid Speciation in Wild Sunflowers.

32. Rieseberg LH and Soltis DE. Phylogenetic consequences of cytoplasmic gene flow in plants. Evolutionary trends in Plants. 1991:5(1):65–84. 10.1007/s00606-006-0485-y

33. Schranz ME, Kantama L, De Jong H, and Mitchell-Olds T. Asexual reproduction in a close relative of Arabidopsis: A genetic investigation of apomixis in Boechera (Brassicaceae). New Phytologist. 2006:171(2):425–438. 10.1111/j.1469-8137.2006.01765.x

34. Slon V, Mafessoni F, Vernot B, de Filippo C, Grote S, Viola B, Hajdinjak M, Peyrégne S, Nagel S, Brown S, et al. The genome of the offspring of a Neanderthal mother and a Denisovan father. Nature. 2018. 10.1038/s41586-018-0455-x

35. Suarez-Gonzalez A, Hefer CA, Christe C, Corea O, Lexer C, Cronk QCB, and Douglas CJ. Genomic and functional approaches reveal a case of adaptive introgression from Populus balsamifera (balsam poplar) in P.trichocarpa (black cottonwood). Mol Ecol. 2016:25(11):2427–2442. 10.1111/mec.13539

36. Taylor SA and Larson EL. Insights from genomes into the evolutionary importance and prevalence of hybridization in nature. Nat Ecol Evol. 2019:3(2):170–177. 10.1038/s41559-018-0777-y

37. Tiley GP, Flouri T, Jiao X, Poelstra JW, Xu B, Zhu T, Rannala B, Yoder AD, and Yang Z. Estimation of species divergence times in presence of cross-species gene flow. Syst Biol. 2023:72(4):820–836. 10.1093/sysbio/syad015

38. Tricou T, Tannier E, and de Vienne DM. Ghost lineages can invalidate or even reverse findings regarding gene flow. PLoS Biol. 2022a:20(9):e3001776. 10.1371/journal.pbio.3001776

39. Tricou T, Tannier E, and De Vienne DM. Ghost Lineages Highly Influence the Interpretation of Introgression Tests. Syst Biol. 2022b:71(5):1147–1158. 10.1093/sysbio/syac011

40. White A, Tolman M, Thames HD, Withers HR, Mason KA, and Transtrum MK. The Limitations of Model-Based Experimental Design and Parameter Estimation in Sloppy Systems. PLoS Comput Biol. 2016:12(12). 10.1371/journal.pcbi.1005227

41. Yakimowski SB and Rieseberg LH. The role of homoploid hybridization in evolution: A century of studies synthesizing genetics and ecology. Am J Bot. 2014:101(8):1247–1258. 10.3732/ajb.1400201

42. Yang Z. The BPP program for species tree estimation and species delimitation.

43. Zheng Y and Janke A. Gene flow analysis method, the D-statistic, is robust in a wide parameter space. BMC Bioinformatics. 2018:19(1):1–19. 10.1186/s12859-017-2002-4

